# Temporal segmentation and ‘look ahead’ simulation: Physical events structure visual perception of intuitive physics

**DOI:** 10.1101/2023.06.14.544968

**Authors:** Tristan S. Yates, Shannon Yasuda, Ilker Yildirim

## Abstract

How we perceive the physical world is not only organized in terms of objects, but also structured in time as sequences of events. This is especially evident in intuitive physics, with temporally bounded dynamics such as falling, occlusion, and bouncing demarcating the continuous flow of sensory inputs. While the spatial structure and attentional consequences of physical objects have been well-studied, much less is known about the temporal structure and attentional consequences of physical events in visual perception. Previous work has recognized physical events as units in the mind, and used pre-segmented object interactions to explore physical representations. However, these studies did not address whether and how perception imposes the kind of temporal structure that carves these physical events to begin with, and the attentional consequences of such segmentation during intuitive physics. Here, we use performance-based tasks to address this gap. In Experiment 1, we find that perception not only spontaneously separates visual input in time into physical events, but also, this segmentation occurs in a nonlinear manner within a few hundred milliseconds at the moment of the event boundary. In Experiment 2, we find that event representations, once formed, use coarse ‘look ahead’ simulations to selectively prioritize those objects that are predictively part of the unfolding dynamics. This rich temporal and predictive structure of physical events, formed during vision, should inform models of intuitive physics.

## Introduction

Against the undelimited influx of sensory inputs arriving at the eyes, our perceptual experiences are richly structured, with objects and events delimiting discrete spatial and temporal entities. Such structure is especially evident in the visual perception of *intuitive physics* — our ability to see, at a glance, and predict how objects move and react to forces. Yet, in many ways, most previous research on intuitive physics has focused on only half of this structure: representations of physical objects. Among others, these studies have illustrated the perception of physical stability of object configurations (Battaglia et al., 2013; Pramod et al., 2022), whether an object is heavier or harder than others (Hamrick et al., 2016; Schwettmann et al., 2019; Guo et al., 2020), and how intuitive physics may be recruited for object perception (Wong et al., 2022). Yet, physical scenes inherently unfold over time, expressing discrete temporal structure in demarcated episodes of object interactions, including collision, bouncing, toppling, falling, and entering and exiting containers. Despite their centrality to our perceptual experience and their importance in the broader cognitive science literature (Michotte, 1963; Gibson, 1954, 1975; Spelke, 1990), the *structure* of physical event representations, as they occur in the visual perception of intuitive physics, remains largely unaddressed.

Consider the simple dynamics of a blue marble rolling on a crowded surface, depicted in Figure 1 as two snapshots over time. When observing the marble, various scene elements ebb and flow into and out of our awareness: We attend to the planks under and across the marble in the earlier moment, and the narrow passage and the green container that the marble will enter in the later moment. It is only natural to describe in *language* a scene of this sort in terms of a sequence of interactions and the objects involved in those interactions (as we just did), but here we ask: Does the *visual system* spontaneously mark such physical event boundaries, even during passive viewing of dynamical scenes? And if so, what are the temporal characteristics of such real-time perceptual segmentation of unfolding dynamics? Moreover, once a segmentation occurs, i.e. between a pair of physical events, what principle(s) guide which objects will be prioritized as part of the current event representation? Answers to these questions will reveal the basic structure of physical events in vision, and the goal of this paper is to address them by introducing new performance-based psychophysical tasks.

**Figure 1.**
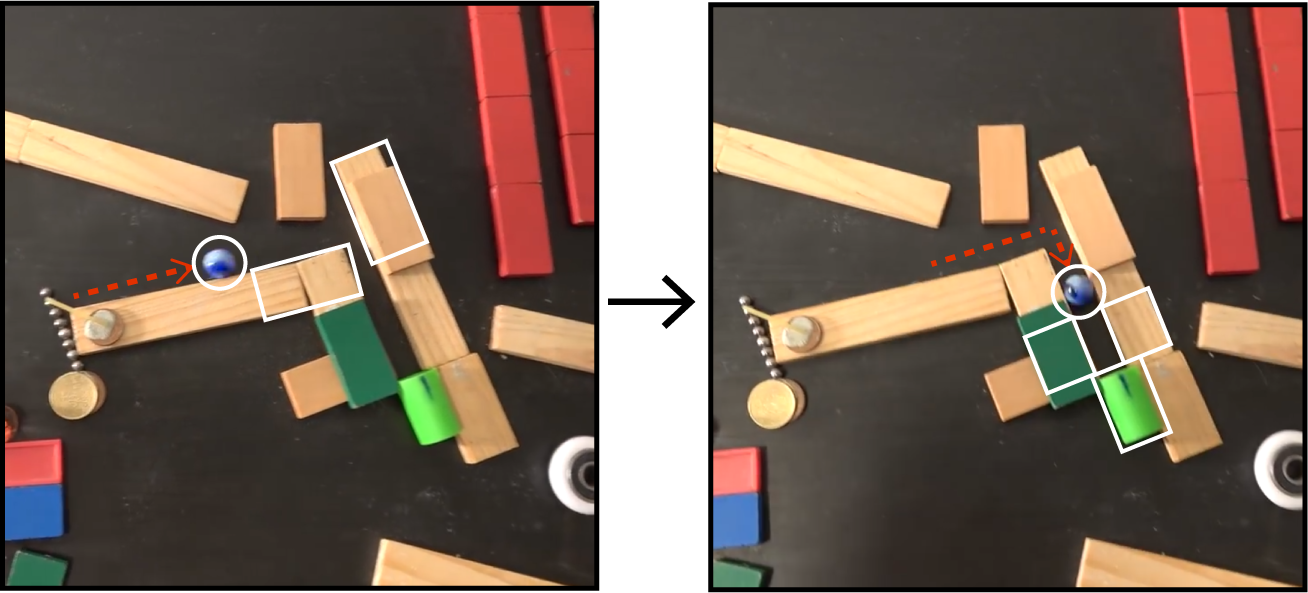
A dynamical scene with a blue marble rolling on a surface and interacting with other objects, shown in two snapshots (source: https://www.youtube.com/watch?v=Hmb0Q0Q_7jo). The dashed red arrow shows the marble’s previous trajectory, and white bounding boxes represent the objects relevant to the unfolding dynamics within each snapshot.

### Physical events as units in the mind

The study of physical events has a long history in cognitive science. Early work examined the perception of (real or apparent; rigid or non-rigid) object motion (Gibson, 1954), as well as how causality is perceived in so-called ‘launching’ events (Michotte, 1963). These examples and others made clear that there is a distinction that can be made between physical events as they occur in the world and physical events as they occur in the mind — in other words, changes that are psychologically meaningful (e.g., the breaking of glass) are not necessarily meaningful in classical physics (since matter is conserved; Gibson, 1975). People seem to have an intuitive understanding of how physical events unfold, even if their judgements do not always adhere to Newtonian physics (McCloskey, 1983). Landmark developmental research demonstrated that expectations about physical events are present early in infancy, suggesting that they may be part of a core system of knowledge (Spelke, 1990; Spelke & Kinzler, 2007).

Much like objects, physical events are only partially determined by the sensory data; rather, they emerge as units in the mind via perceptual processing, and provide a natural interface for attentional selection. Yet, previous work in intuitive physics either does not address physical events, or considers physical events as a given, as pre-segmented interactions in isolation: one object moves, one object falls, or one object is occluded. This is in contrast to how we normally encounter dynamic scenes, where an object in motion may participate in several different types of events, recruiting, at each event, a different *causal context* — varying subsets of all visible objects and surfaces relevant to ongoing dynamics. Even though using *isolated* physical events has led to key insights about physical representations (e.g., Gerstenberg et al., 2017; Little & Firestone, 2021; Karakose-Akbiyik, Caramazza, & Wurm, 2023), a full understanding of intuitive physics requires investigating how its temporal structure, i.e., sequences of physical events, arises during the processing of continuous visual input, and how physical events impact attentional prioritization of the visible objects in a scene, particularly at the time scales relevant to fast perceptual processing. Therefore, to make progress in intuitive physics, it is essential to study the *formation* and *contents* of physical events during continuous visual processing.

### Uncovering physical event structure in perception: The current study

In the broader event cognition literature, two main paradigms have been used to measure event segmentation: the unitization task (Newtson & Engquist, 1976; Radvansky & Zacks, 2017; Zacks et al., 2007; Zacks, 2020) and the dwell-time paradigm (Hard et al., 2011; Zheng et al., 2020). The unitization task, as an explicit judgment task, is not well-suited for revealing the spontaneous, automatic construction of event representations within visual perception. The dwell-time paradigm, on the other hand, while relatively more implicit, relies on participants advancing videos at their own pace, and is therefore not only insufficiently granular for fast timescale events like physical events, but also interferes with relevant aspects of the stimuli (e.g., the velocity of objects). A different but related method comes from the domain of audition and music cognition. Repp (1992, 1998) showed that when participants listened to a musical piece where certain notes were extended in time relative to the original musical score, participants were less likely to detect these changes at the transitions (i.e., boundaries) between phrases. As we show in this article, this method suggests a paradigm for visual perception that is more objective via its performance-based manner, allowing us to determine if boundaries between physical events are spontaneously formed and segmented during seeing (see also Ji & Papafragou, 2022).

In perception, segmenting time is not the goal in itself, but a means to a more tractable setting for selective processing of otherwise complex, dynamic visual environments. For example, event representations guide which aspects of an object are prioritized: Subtle changes made to the relevant dimension of an object (height vs. width) with respect to the given event category (occlusion vs. containment) are differentially more detectable (Strickland & Scholl, 2015). Additionally, seeing an actor’s reaction to invisible surfaces or objects implies their physical presence, influencing participants’ responses to congruent or incongruent probes (Little and Firestone, 2021). How do events guide such selective processing in perception? One principle advanced and strongly supported in the event cognition literature is the idea of *prediction* (Reynolds et al., 2007; Zacks et al., 2007, 2011): Event models should compose of elements that will facilitate prediction during the epoch of that event, with boundaries between events signaled by a decreased ability to predict what will happen next, known as *prediction errors*. Applied to physical events, this principle suggests a selective processing mechanism that should take into account the simulated trajectories of objects (i.e., it should ‘look ahead’ in time) to prioritize those objects that will impact dynamics within the locality of the current event.

In the current project, we investigate the basic structure of physical event representations within continuous perception by addressing (1) the time course of how a new event is formed — i.e., when boundaries are imposed on continuous input — and (2) what determines the contents of an event representation. Experiment 1 generalizes the performance-based tasks developed in the domain of music cognition (Repp, 1992, 1998) to the domain of seeing dynamical scenes. Based on prior work, we predict that participants’ detection accuracy for transient slow-downs that occur during videos of multiple physical events will decrease at event boundaries relative to non-boundary intervals (although it is also possible that detection accuracy may increase at boundaries; Baker & Levin, 2015). This task allows us to investigate the sub-second time course of event segmentation in terms of how detection accuracies rise and fall, and the symmetry of this pattern along the event boundary. In Experiment 2, we predict that participants will be better at detecting ‘simulation’ probes — briefly presented letters on stationary objects in dynamical scenes — if they appear on objects that are along the simulated trajectory of the current event, relative to spatially controlled counterparts. This task allows us to test the hypothesis that physical event representations that are formed spontaneously during vision guide selective processing via prediction, and whether this is modulated by event boundaries. Critically, in both experiments, participants are not informed of physical events and are simply tasked with paying attention to either temporal (Experiment 1) or simulation (Experiment 2) probes. Thus, our study design should tap into automatic, visual processes and help reveal the structure of physical event representations as they are formed during vision.

#### Experiment 1

In Experiment 1, we examined how and when physical events are spontaneously segmented as part of visual processing. To reveal the time course of how physical event segmentation occurs at sub-second resolution, we showed participants videos of multiple physical events and asked them to indicate if and when during the video they noticed a temporary slowing of the video.

## Methods

### Participants

Per our pre-registration, we recruited 110 participants from Prolific, such that there were 11 groups of 10 participants who saw the same set of videos. We excluded 1 participant from our subject-wise analyses for not having at least half of the trials usable, and 1 additional participant for not having enough event boundary trials usable (see below for pre-registered trial exclusion criteria). This number of participants does not include data from an additional 29 participants who started the task but did not complete all of the test trials. Participants (N=50 female, N=58 male, N=2 unknown) were between 20 and 37 years of age (Mean age = 28.05 years) and were majority white (N=80 White, N=7 Black, N=9 Asian, N=9 multiple ethnicities, N=2 other, N=3 unknown) from the United States or United Kingdom (N=29 US, N=79 UK, N=3 unknown).

### Stimuli

We created multi-event videos using a combination of Python (Van Rossum & Drake, 2009) and Blender (Community, 2018) software, inspired by recent work on causal graph structures in physics representations (Roussel et al., 2019). Specifically, we modified the code base created by Roussel and colleagues to compose videos where objects interacted with one another across multiple different physically-plausible events. We focused our research on a handful of event types that were simple and easily identifiable by human participants: collision (two objects collide), occlusion (one object goes behind another or becomes visible again), containment (one objects enters another object), falling (one object falls from a ramp), and toppling (one unstable object hits the ground; see Figure 2A). For the current experiment, we focused on the former 4 event types, as they could easily be strung together into every possible two-event combination (e.g., an object falls and then collides with another object; an object is occluded and then contained by another object, etc.). We created two unique examples for each two-event combination for a total of 32 ‘base’ videos (Mean video duration: 9.75 seconds; Range: 6.23 – 14.3 seconds). Additionally, we added a 3-second countdown to the start of each video so that participants could prepare for the video onset.

**Figure 2.**
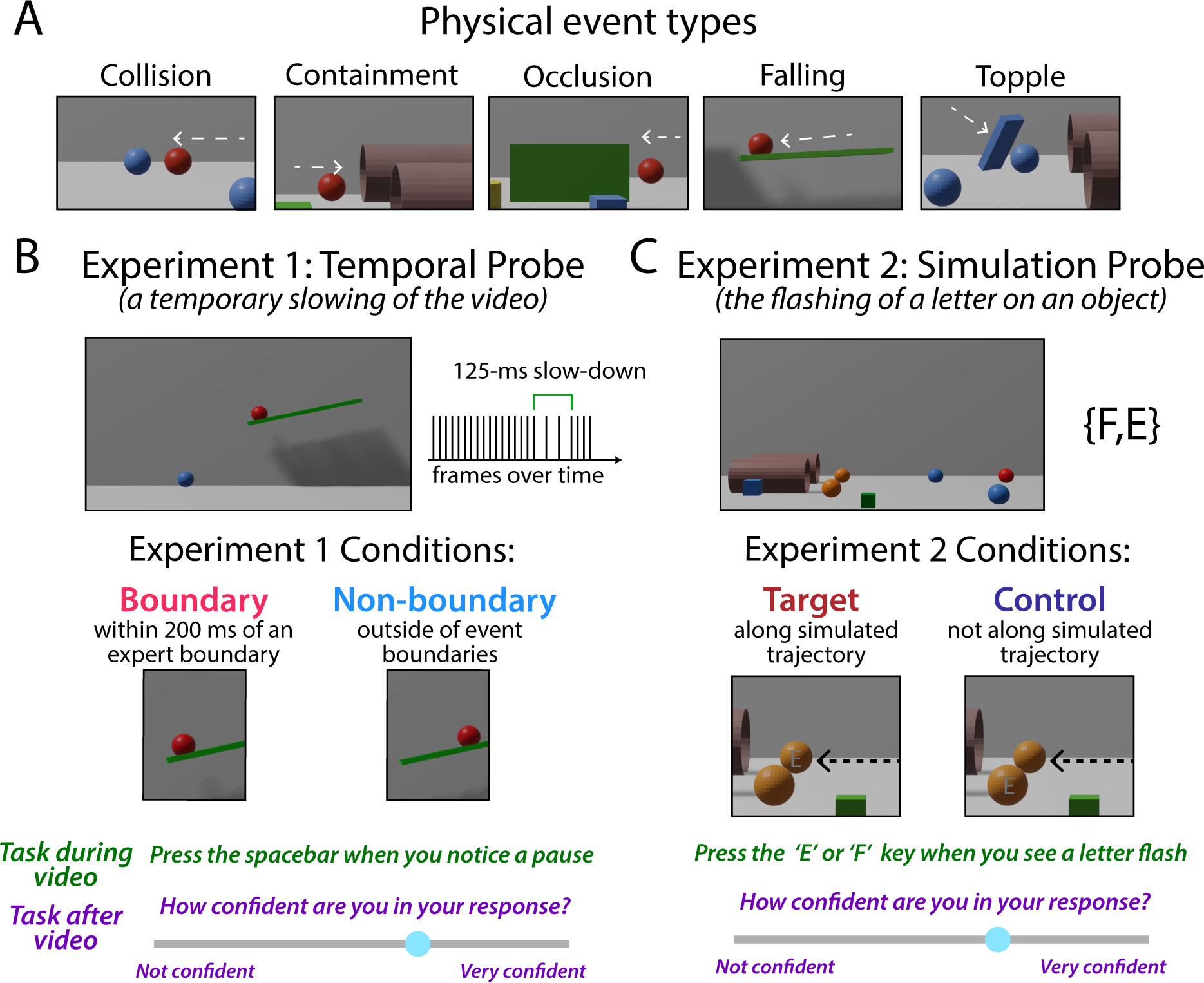
Design of the two experiments presented in this paper. (A) In both experiments, participants saw videos that consisted of a sequence of two or more different physical event types: collision, containment, occlusion, falling, and topple. Dashed arrows indicate the motion of the target object – the object that is set in motion initially in each scene – and were not shown during the actual experiment. (B) In Experiment 1, participants watched videos with or without temporal probes (temporary slowing of the video for 125 milliseconds) that occurred either at expert-determined event boundaries (e.g., the onset of a ball falling off of a ramp) or at other points in the video. Participants were tasked with pressing the spacebar when they noticed a temporal probe in the video and reporting their confidence at the end of the video. (C) In Experiment 2, participants watched videos with or without simulation probes (briefly presented letters on stationary objects) that occurred either on objects that were on the simulated trajectory of the ongoing event (e.g., the orange ball that the red ball will hit; indicated with the dashed black arrows) or matched objects that were outside of the ongoing event. Participants were tasked with pressing either the ‘E’ or ‘F’ key when they saw the letter flash on the screen, and reporting their confidence at the end of the video.

Physical event boundaries were defined by two expert coders (authors) who resolved disagreements by discussion. The criteria for determining physical event boundaries were subjective, but coders aimed to identify the onsets and offsets of every physical event shown in the scene. Each video was determined to have 2 or 3 event boundaries that separated different physical event types (e.g., the end of an occlusion event prior to a collision event). These event boundaries were used to relate to temporal probe detection accuracy.

### Temporal probe task

For the temporal probe task, participants were asked to quickly and accurately respond to a 125-millisecond slowing of the video frame rate by pressing the spacebar. Videos were shown centrally on the screen with a reminder at the top of the page: ‘Press the spacebar when a pause occurs.’ The border around the video flashed red when it registered a spacebar response; otherwise, the border was black. After each video finished, participants rated their confidence on a sliding scale from ‘Not very confident’ to ‘Very confident’ (Figure 2B). Participants could then press the next button to start the next trial. Progress through the experiment was shown with a trial counter on the bottom of the screen.

Temporal probes could occur at any 200-millisecond time point from the start of a video, as long as there was motion on the screen (e.g., probes did not occur when a ball was occluded). Across the 32 base videos, there were many possible temporal probe locations (Mean: 42, range: 26 – 64), but we only sampled 22 of these probe locations for each video. To do this, we created 11 different sets of videos (each containing 96 videos), and collected data from 10 participants for each set. All participants completed a total of 96 trials, where 64 videos contained a temporal probe, and 32 did not. Some temporal probes (average of 4.85 per participant) occurred at event boundaries (e.g., as a ball was going to fall off of a ramp), while the majority of temporal probes (average of 59.15 per participant) occurred at timepoints within events (e.g., a ball further up the ramp; Figure 2C). The same base videos were shown three times throughout the experiment — twice with a temporal probe and once without it (as a filler). We only analyze the trials in which a temporal probe was presented. Videos were shuffled and presented in a random order for each participant. Per our pre-registration, individual trials were excluded if participants pressed the spacebar more than once during a video, or if participants responded outside of the acceptable response time window (within 300 and 2500 ms of the probe).

At the start of the experiment, participants were asked to scale a bounding rectangle to the size of a credit card, to ensure that videos were approximately the same size across participants. They then were shown the experiment instructions and completed two practice trials. On the first practice trial, participants were shown a video of a ball colliding with another ball and then a third ball, and told there would not be a temporal probe in this video. On the second practice trial, the same video was shown, but now there was a 125-millisecond slowing of the video as the ball rolled towards the first ball it was going to collide with. After the practice, participants completed a two-question quiz on the instructions to test for comprehension. If either question was answered incorrectly, the instructions and practice trials were repeated, and participants only proceeded to the main experimental trials once they obtained 100% on the comprehension test. After completing the experimental trials, participants were asked a series of debriefing questions about task difficulty, engagement, and any feedback they had about the task. The code used to run this experiment was built using psiTurk, a platform for running behavioral experiments online (Eargle et al., 2020; Gureckis et al., 2016).

### Event segmentation task

To supplement the expert-determined boundaries and to test whether naive participants’ explicit boundary judgments relate to detection accuracy on the temporal probe experiment, we additionally collected event segmentation data from a separate set of Prolific participants. We had 20 new participants complete this explicit segmentation task, which does not include 6 participants who did not finish the task. Demographics of these participants were similar to the temporal probe task: Participants (N=14 female, N=6 male) were between 21 and 37 years of age (Mean age = 29.20 years) and were majority white (N=16 White, N=3 Asian, N=1 other) from the United States or United Kingdom (N=1 US, N=19 UK). For each of the 32 base videos, participants were first shown the video in its entirety without the ability to pause or rewind. Then, participants were able to scroll through the video and press a key to indicate that an event boundary occurred at a given time window. Participants were allowed to denote multiple event boundaries and were free to change their answers by pressing the same key again when they landed on the previously-determined event boundary frame. Once participants were satisfied, they could proceed to the next trial. Example aggregated responses on several typical base videos can be seen in Figure S1. As can be appreciated, while there was some variability in where participants decided an event boundary occurred, the peaks in responses aligned well with expert-determined boundaries.

## Results

### Detection accuracy: People are worse at detecting probes that occur at event boundaries

For our main analysis, we asked whether detection accuracy for temporal probes placed at expert-determined boundaries (i.e., between collision, containment, falling, and occlusion events) was lower than temporal probes placed at non-boundary timepoints. In this and all following analyses, we only consider the trials in which participants responded once within the detection window (average 1.26% of trials per participant dropped for multiple key presses; average 8.64% of trials per participant dropped for out-of-range response times). Probe detection accuracy was averaged across boundary and non-boundary probe trials within each participant, and a two-sided paired *t*-test was used to determine significance. As depicted in Figure 3A, participants’ detection accuracy for temporal probes that occurred at event boundaries (*M* = 33.20%) was lower than that of non-event boundaries (*M* = 61.06%; *t*(107) = -11.36, *p* < .001) — a strikingly strong effect (Cohen’s *d* = 1.10), that was evident in 96 of the 108 participants we analyzed (Figure 3B). This was also true in our pre-registered analysis collapsing across participants (instead of trial types within participant): using Fisher’s exact test, we found that temporal probes that occurred at event boundaries (36.32% of 479 boundary probes detected) were less likely to be detected compared to non-event boundaries (62.67% of 5760 non-boundary probes detected; two-sided test, *p* < .001). Thus, participants readily experience the boundaries between physical events, as evidenced by their difficulty in recognizing changes in the speed of a video at event boundaries.

**Figure 3.**
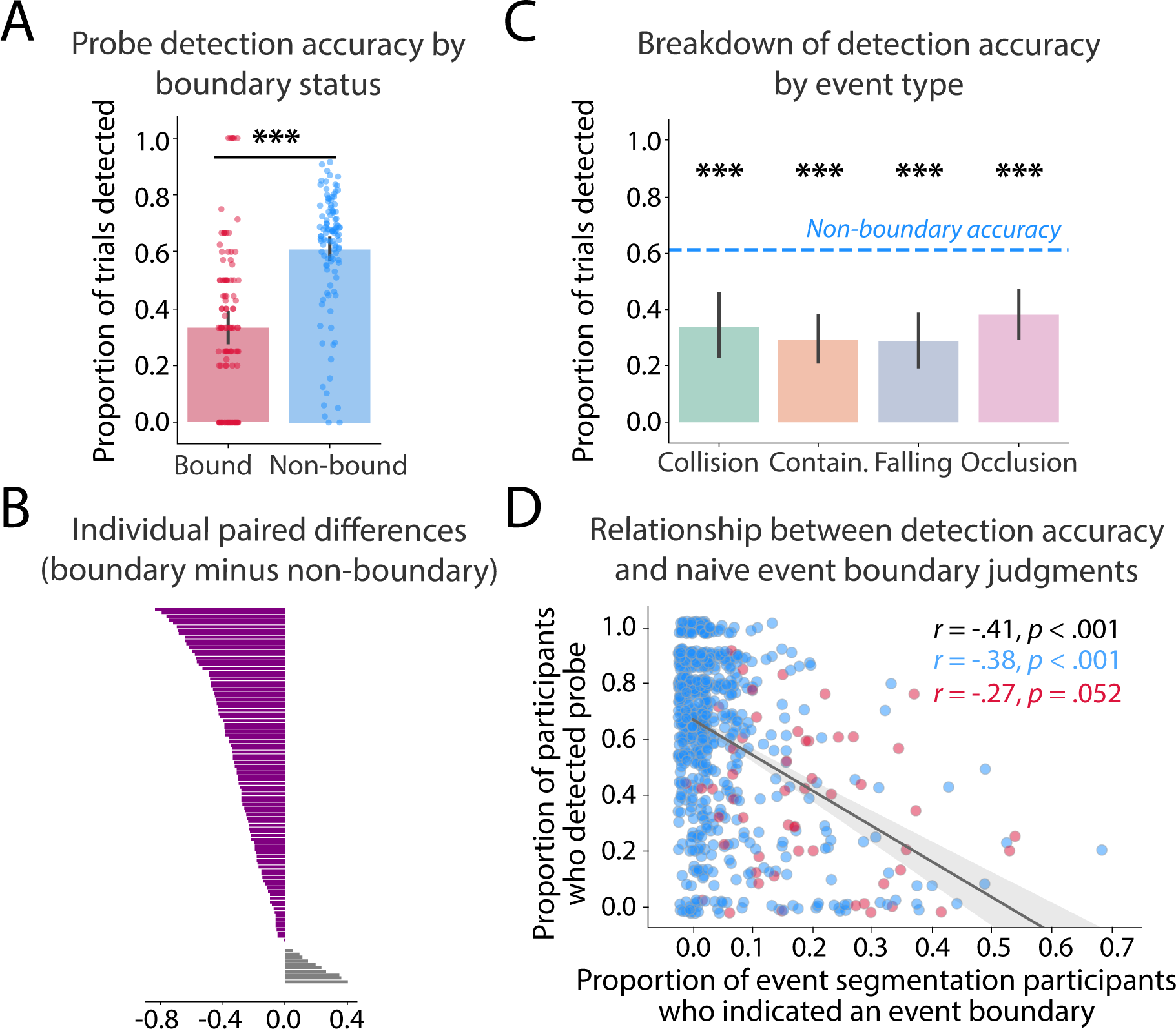
Temporal probe detection accuracy from Experiment 1, averaging the boundary and non-boundary responses within participants. (A) Temporal probes were significantly less likely to be detected when they occurred at expert-determined event boundaries compared to non-boundaries. Dots represent individual participant averages, and error bars reflect the 95% confidence interval derived from bootstrap resampling of the mean. (B) Differences at the individual participant level, sorted by the size of the effect. The expected difference (boundary accuracy less than non-boundary accuracy) was found in 96 out of the 108 participants tested. (C) Exploratory breakdown of temporal probe detection accuracy for the four different event types. Average participant accuracy on non-boundary trials is marked with a dashed line. Because the number of event type boundaries that were tested varied by participant group, subsets of participants are included in this analysis per event type (collision: N=68; containment: N=94; falling: N=75; occlusion: N=76). (D) The proportion of participants who detected a temporal probe was significantly negatively related to the proportion of event segmentation participants who indicated an event boundary in the 200 ms window around that time point when considering all possible probes, and when specifically looking at non-boundary probe trials (blue dots), but only marginally so when considering boundary probe trials (red dots). Dots represent averages across participants for different probe trials. The shaded area reflects the 95% confidence interval derived from bootstrap resampling of the correlation. *** *p* < .001.

As an exploratory follow-up, we investigated probe detection accuracy in each event type separately (Figure 3C). Because different participants saw different subsets of physical events, varying numbers of participants contributed data to this analysis. Across all event types, detection accuracy was substantially decreased relative to non-boundary probe detection accuracy (collision: *M* = 33.82%; *t*(67) = -5.34, *p* < .001; containment: *M* = 29.26%; *t*(93) = - 6.95, *p* < .001; falling: *M* = 28.60%; *t*(74) = -7.40, *p* < .001; occlusion: *M* = 38.16%; *t*(75) = - 5.13, *p* < .001). This was also true when we performed our pre-registered analysis of collapsing across participants (collision: 42.86%; *p* < .001; containment: 37.13%, *p* < .001; falling: 26.77%, *p* < .001; occlusion: 40.86%, *p* < .001). This suggests that the overall lower detection accuracy we see is not being driven by a particular event type, but may be a shared property of physical events.

In addition to investigating probe detection accuracy at expert-determined boundaries, we asked whether probe detection accuracy tracks naive participants’ agreement about the placement of event boundaries. To test this, we averaged probe detection accuracy across participants for each possible temporal probe (704 unique videos) — on average, this value represented detection accuracy aggregated from 8.9 participants per temporal probe. Then, we calculated for each of the 32 base videos the proportion of event segmentation participants who indicated an event boundary had occurred within 200 ms time windows. Finally, we correlated average probe detection accuracy from Experiment 1 with the proportion of participants who indicated an event boundary within the window of time of the corresponding temporal probe. When considering all trials (including both the boundary and non-boundary trial types), we found a significant negative relationship between probe detection accuracy and event boundary agreement (*r* = -.41, *p* < .001), indicating that the more people agreed that an event boundary occurred, the less likely a temporal probe would be detected at that time point. This relationship persists when we only consider non-boundary probe trials (*r* = -.38, *p* < .001), and is marginally significant when we only consider boundary probe trials (*r* = -.27, *p* = .052). These results illustrate the robustness of our categorical analysis and further establish that event boundaries are experienced during spontaneous visual processing.

Our results thus far have not taken into consideration an important potential confound: that is, at event boundaries, we may expect to see more changes in the low-level pixel values of the video. This means that we may not be picking up on event boundary representations *per se*, but instead the impact of image changes on temporal probe detection accuracy. To address this, we first calculated the amount of pixel change that occurred during temporal probes separately for boundary and non-boundary probe trials. Specifically, we used OpenCV (Bradski, 2000) to average the difference in pixel values between adjacent frames for each frame during the temporal probe. First, we found that the amount of pixel change that occurred for temporal probes at event boundaries (*M* = 0.06) was not different from that of non-boundaries (*M* = 0.05; *t*(703)=1.36, *p* = .173). Second, we investigated whether boundary status still had a significant impact on temporal probe detection accuracy when we include pixel change as a predictor. In a linear mixed effects model with participant as a random effect, we found that pixel change did not predict temporal probe detection accuracy (*β* = 0.13, *p* = .812), while boundary status did (*β* = -1.09, *p* < .001; Table 1). Therefore, our results are inconsistent with a purely low-level account of pixel change.

**Table 1.** Results from a linear mixed effects model predicting probe detection (1 for detected, 0 for not detected) from the amount of pixel change at the probe and the boundary condition (1 for boundary, 0 for non-boundary), using data from all trials across participants (with participant as a random effect). The equation used in the model is shown as the table heading. The beta parameter, standard error, *t*-statistic, and *p*-values are given for each of the predictors.

|  | Beta estimate | Std. error | *t*-statistic | *p*-value |
| --- | --- | --- | --- | --- |
| Intercept | 0.613 | 0.020 | 30.695 | < .001 |
| Condition (boundary) | -0.270 | 0.022 | -12.491 | < .001 |
| Pixel change | 0.015 | 0.121 | 0.123 | .812 |
| Participant random effect | 0.036 | 0.012 | - | - |

### The time course of event segmentation: Temporal symmetry in probe detection relative to event boundaries

So far, the results support our hypothesis that participants segment the continuous flow of sensory inputs according to physical events, as evidenced by their decreased ability to detect temporal probes at expert-determined event boundaries as well as boundaries empirically determined by naive participants.

Critically, beyond the basic contrast of boundary versus non-boundary trials, our experimental design allows us to investigate the time course of how a new event is segmented in visual perception. We analyzed how probe detection accuracy changes as a function of time — i.e., when during the video the probe occurred relative to the moment of the nearest event boundary. To do so, we first collapsed the detection accuracy across trials, resulting in between 190 and 483 responses over the time interval of -1000 milliseconds to 1000 milliseconds centered at the event boundary at increments of 200 milliseconds. Second, we calculated confidence intervals for detection accuracy at each temporal distance to the event boundary by resampling the corresponding trials 1000 times with replacement. Finally, we characterized the rate of change in detection accuracy using a sigmoid function, instead of a linear fit, based on our qualitative observations that the way detection accuracy changed as a function of time was often non-linear. We separately fit sigmoid functions pre-event boundary (between -1000 ms and 0 ms) and post-event boundary (between 0 ms and 1000 ms) using SciPy curve fitting functions (Virtanen et al., 2020). Specifically, we conducted 1000 bootstrap resamples of the average detection accuracy for each window of time, fitting a sigmoid function and saving the parameter values on each iteration. We then performed a two-tailed direct bootstrap hypothesis test on the resulting parameter estimates for the logistic growth/decay rate to assess whether the rates of change were symmetric or significantly different pre- and post-boundary.

As shown in Figure 4A, there was a drop-off in average temporal probe detection accuracy when approaching the event boundary, followed by a return to baseline accuracy following the event boundary. Roughly centered at the event boundary, there is approximately one full second period (from -500 milliseconds to 500 milliseconds relative to the boundary) in which accuracy drops by a large amount and recovers that back. Indeed, this was confirmed in our analysis of the resampled sigmoid slope values: the rate of logistic decay pre-boundary (log(*β*) = 2.86) was not significantly steeper than the rate of logistic growth post-boundary (log(*β*) = 1.23; *p* = .168; Figure 4B). However, a bimodal distribution for pre-boundary rates of logistic decay was evident, with some values being in the same range of the post-boundary rate of logistic growth, and some seemingly higher. We next investigated whether this bimodal distribution may be due to differences across different types of event boundaries. Indeed, we found that the rates of logistic growth and decay in detection accuracy somewhat varied by event type (Figure 4C). Specifically, consistent with the overall trend, pre-boundary rates of change were not significantly different from post-boundary rates of change for containment (log(*β*) = 1.50 vs. log(*β*) = 0.77; *p* = .576) or falling events (log(*β*) = 3.66 vs. log(*β*) = 3.32; *p* = .640). However, for the remaining event types, it is evident that there is a quick drop-off in detection accuracy pre-boundary, followed by a more gradual pickup post-boundary; for collision, pre-boundary rates of change were significantly higher/steeper than post-boundary rates of change (log(*β*) = 3.63 vs. log(*β*) = 0.78; *p* = .048), and the same was true for occlusion (log(*β*) = 3.34 vs. log(*β*) = 0.53; *p* = .014). These results indicate that while there is an overall decrease in temporal probe detection accuracy at event boundaries, how soon this detection accuracy decreases and then returns to baseline accuracy level seems to depend on the particular event types. These exploratory results indicate that there may be differences in how event models come online during perception.

**Figure 4.**
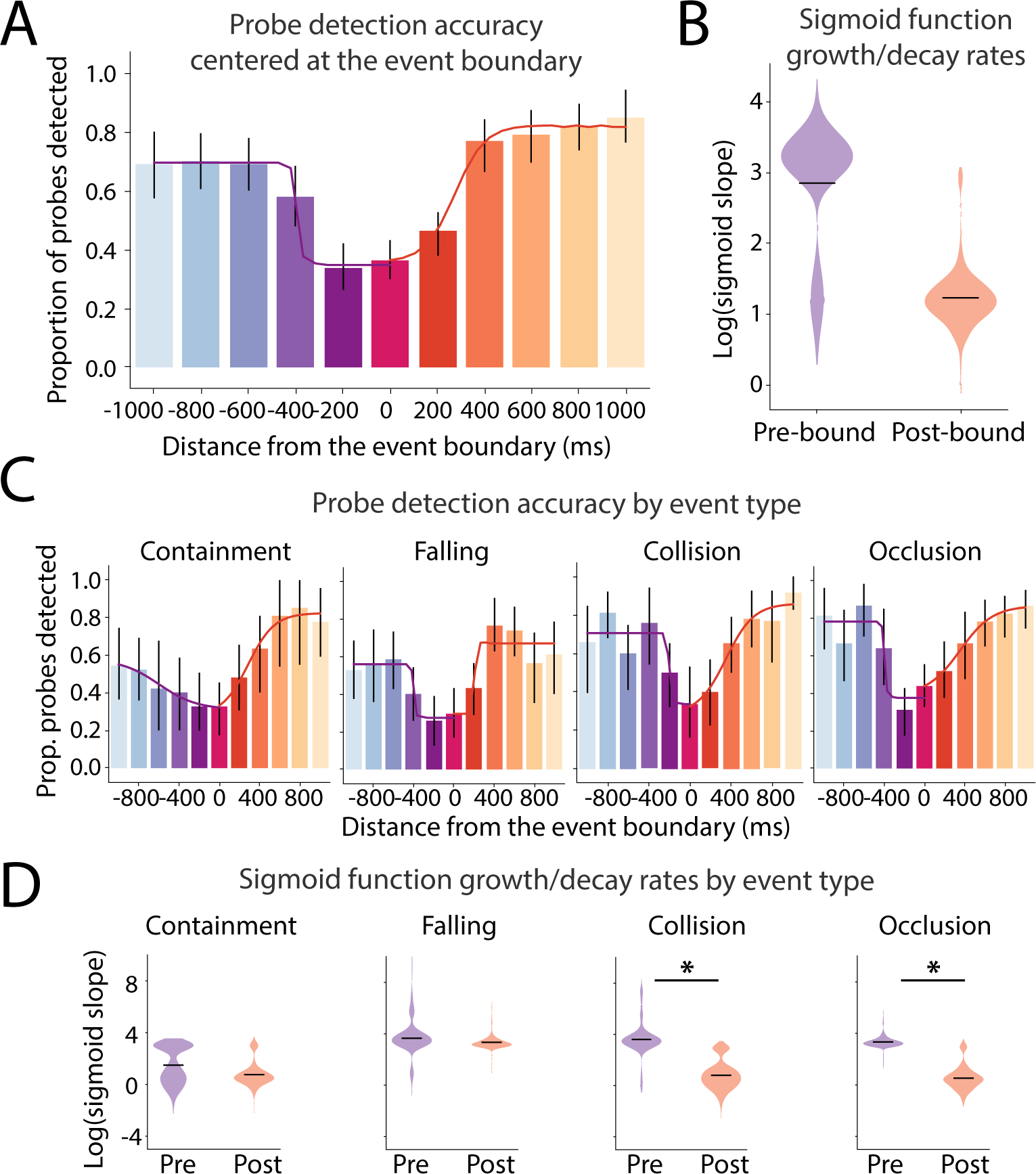
Rates of change in the average probe detection accuracy approaching and receding from expert-determined event boundaries. (A) Detection accuracy rapidly declines prior to event boundaries, with the lowest detection accuracy occurring 200 ms before the boundary, and then increases following the event boundary. Sigmoid functions quantify such change in detection accuracy, and the best fits to the average data are displayed as lines pre-(purple) and post-boundary (orange). (B) Sigmoid model slopes of the relationship between detection accuracy and temporal distance from the event boundary, calculated separately for pre-boundary and post-boundary. Log-transformed slopes are shown based on 1000 bootstrap resamples. The average sigmoid model slope was not significantly different pre- and post-boundary. (C) Detection accuracy and distance plotted separately for each of the four event types. (D) The sigmoid model slopes after log transformation for 1000 bootstrap resamples plotted separately for each of the four event types. * *p* < 0.05.

### Confidence and response time also reflect event boundaries

The focus of our research was on temporal probe detection accuracy, but we can also ask whether participants’ confidence and response times were impacted by whether temporal probes were placed at boundary vs. non-boundary timepoints. First, we investigated probe detection confidence averaged across boundary and non-boundary probe trials within each participant using a two-sided paired *t*-test. As depicted in Figure S2A, participants’ confidence in their performance at detecting temporal probes at event boundaries (*M* = 69.46) was lower than their confidence for detecting temporal probes at non-event boundaries (*M* = 75.44; *t*(107) = -5.52, *p* < .001). This is consistent with our main detection accuracy findings, which suggest that detecting temporal probes at event boundaries is more difficult. Importantly, although the amount of pixel change that occurred during a probe was related to participants’ confidence, there was still a significant effect of event boundary status (Table S1). There was also a significant relationship between confidence and event boundary agreement across our event segmentation participants for all probe trials (*r* = -.20, *p* < .001) and when only considering non-boundary probe trials (*r* = -.14, *p* < .001), but not when only considering boundary probe trials (*r* = -.13, *p* = .340; Figure S2B).

Finally, we asked whether participants were slower to respond to temporal probes when they occurred at event boundaries. Similar to our confidence results, we found that event boundary status predicted response time such that even if people detected a probe, they were in fact slower to respond when that probe occurred at an event boundary. This result is based on a multiple regression model that took into account pixel change (Table S2). We find broadly consistent (though weaker) results in participant-level and continuous analyses of response time data (Figure S2C, D).

#### Experiment 2

Experiment 1 demonstrated how and when visual input is demarcated into discrete events during perception. However, this is only important insofar as it helps guide attention and other downstream processing. In Experiment 2, we tested whether a form of ‘look ahead’ simulations guide which objects are prioritized as part of a physical event representation. We showed participants videos consisting of multiple physical events in a sequence and asked them to indicate if and when during the video they noticed the flashing of a letter on one of the objects in the scene.

## Methods

### Participants

We recruited undergraduate students from the Psychology Subject Pool to complete an online version of this task. Our goal was to recruit 20 total participants in 2 groups of 10 participants each, with participants in the same group seeing the same combination of videos. However, due to the asynchronous nature of online recruitment, we ended up collecting 28 total participants (13 saw one set of videos, and 15 saw the other set). All participants were included in the study for having more than half of the trials usable (described more below). Demographic information of the participants is unavailable, but Psychology Subject Pool participants tend to be college-aged (18 to 22 years old) and balanced in terms of gender.

### Stimuli

We created new multi-event videos again using a combination of Python and Blender software, as in Experiment 1. This time, we used all five event types (collision, occlusion, containment, falling, and toppling) and strung them together into three-event videos. We also added numerous distracting objects to each scene in order to test for attentional allocation. We created a total of 32 ‘base’ videos that were between 3 and 6 seconds long (Mean video duration: 4.39 seconds).

From these base videos, we created probe videos using Adobe Premiere Pro software (Adobe Inc., 2019). Specifically, for each video, we first marked the transitions between physical events, and then determined whether in the time shortly before that event boundary there was (1) a target object present that would be on the simulated trajectory of the ongoing physical event (e.g., a ball that is about to be collided) and (2) a control object present that was of similar size, shape, color, and location as the target object but not involved in the physical event. Most videos had either one or two possible event boundaries that could be probed, with an average of 1.8 events.

Then, for these events, we created two videos: one in which a gray letter flashed on the target object and one in which a gray letter flashed on the control object (Figure 2C). The letters were always placed in similar locations on the control and target objects, and lasted for 20 milliseconds before disappearing. Videos were randomly assigned to either having an ‘E’ or ‘F’ flash on the screen, but were made consistent across participants. On average, the offset of the letter flash occurred just 35.60 milliseconds before event boundaries, although there was some variability in when these probed occurred, which we later explore.

### Simulation probe task

For the simulation probe task, participants were asked to quickly and accurately respond to the 20-millisecond letter flash by pressing the key that corresponded to the flashing letter (either an ‘E’ or an ‘F’). Right before each trial, participants saw a reminder at the top of the page: ‘Pay attention for an ‘E’ or ‘F’ flashing on the screen.’ In pilot testing, we determined that the size and duration of the simulation probe made it very difficult to detect. Thus, each video was shown in full-screen mode during the actual task. The border around the video flashed red when it registered a response; otherwise, the border was black. After each video finished, participants rated their confidence on a sliding scale from ‘Not very confident’ to ‘Very confident’. Participants could then press the next button to start the next trial. Progress through the experiment was shown with a trial counter on the bottom of the screen.

As mentioned above, simulation probes could occur either on target or control objects shortly before the boundary between physical events. We created 2 different sets of videos, where the same event boundary was only ever tested in either the target or control condition. In one set, 32 targets were probed while 26 control objects were probed; in the other set, the opposite was true. In half of these trials, the letter ‘E’ appeared and in the other half, the letter ‘F’ appeared. All participants also saw 6 videos in which no probe occurred, which were randomly chosen for each participant; however, we only considered trials in which a probe was presented for our analyses. This meant in total, participants completed 64 trials, which were presented in a randomized order. Similar to Experiment 1, individual trials were excluded if participants pressed either the ‘E’ or ‘F’ key more than once during a video, if they responded with the wrong key press, or if they responded outside of the acceptable response time window (within 300 and 2500 ms of the probe).

At the start of the experiment, participants were shown the experiment instructions and completed two practice trials. On the first practice trial, participants were shown a video of a ball rolling out of a tube then hitting a block that topples and collides with another ball. They were told there would not be a simulation probe in this video. On the second practice trial, the same video was shown, but now the letter ‘E’ flashed on the block that would topple. (Participants were shown a screenshot of the letter on the block prior to the practice trial to give them an idea of what to look for; this was never shown for the actual test trials.) After the practice, participants completed a two-question quiz on the instructions to test for comprehension. If either question was answered incorrectly, the instructions and practice trials were repeated, and participants only proceeded to the main experimental trials once they obtained 100% on the comprehension test. After completing the experimental trials, participants were asked a series of debriefing questions about task difficulty, engagement, and any feedback they had about the task. The code used to run this experiment was built using psiTurk (Eargle et al., 2020; Gureckis et al., 2016).

## Results

### Detection accuracy: People are more accurate at detecting probes placed on objects that lie along the simulated trajectory of the ongoing physical event

For our main analysis, we asked whether detection accuracy for simulation probes placed on target objects (i.e., objects that are stationary at the moment when the simulation probe flashes but otherwise lie on the simulated trajectory of the current event) would be *higher* than detection accuracy for simulation probes placed on control objects (i.e., objects that are also stationary at the moment when the simulation probe flashes but lie outside of the simulated trajectory of the current event). The rationale for this hypothesis was that attention would be allocated via a form of ‘look ahead’ simulations toward those objects that lie within the unfolding local causal context of the current physical event (though again, critically, are not actively in motion). To test this, we averaged probe detection accuracy across target and control probe trials within each participant, and used a two-sided paired *t*-test to determine significance. In this and all following analyses, we only consider the trials in which participants responded correctly once within the detection window (average 11.38% of trials per participant dropped for incorrect key press [e.g., ‘F’ when the probe was ‘E’]; average 1.00% of trials per participant dropped for multiple key presses; average 4.41% of trials per participant dropped for out-of-range response times). As depicted in Figure 5A, participants’ detection accuracy for simulation probes that occurred on target objects (*M* = 49.31%) was significantly higher than that of control objects (*M* = 41.71%; *t*(27) = 3.54, *p* = .001, Cohen’s d = 0.68). This effect was not driven by just a few subjects; instead it was observed in 23 out of 28 participants (Figure 5B). These results suggest that participants are indeed selectively processing objects that are along the simulated trajectory of the current physical event.

**Figure 5.**
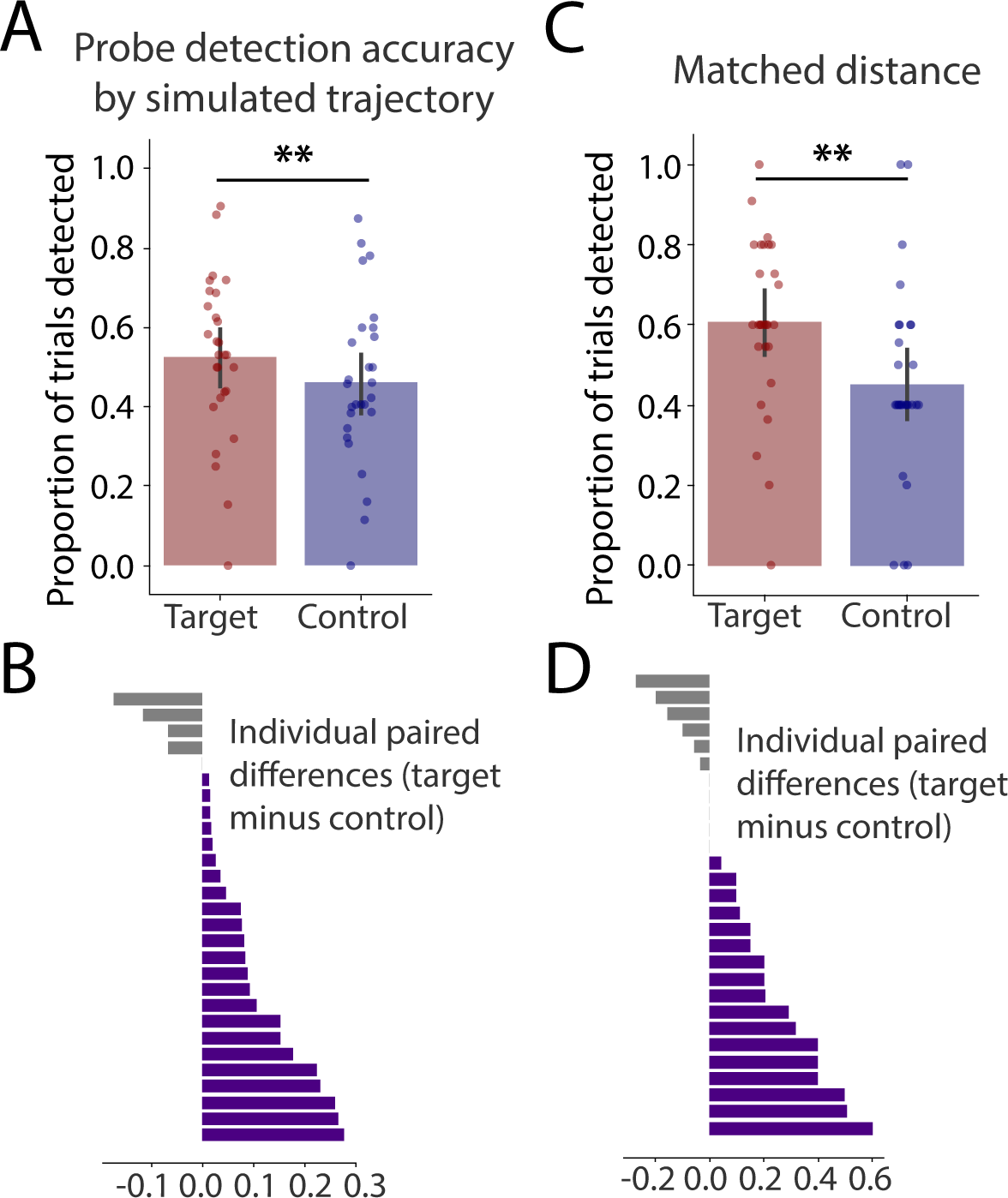
Simulation probe detection accuracy in Experiment 2, averaging the target and control object trials within participants. (A) Simulation probes were significantly more likely to be detected when they appeared on target objects (i.e., objects within the current physical event) compared to control objects. Dots represent individual participant averages, and error bars reflect the 95% confidence interval derived from bootstrap resampling of the mean. (B) Differences at the individual participant level, sorted by the size of the effect. The expected difference (target accuracy greater than control accuracy) was found in 23 out of the 28 participants tested. (C) When matching for the Euclidean distance between the probed objects and the agent object, simulation probes were still significantly more likely to be detected if they appeared on target compared to control objects. (D) Differences at the individual participant level after matching for difference, with the expected difference found in 17 out of the 28 participants tested. ** *p* < .01.

Could these results simply reflect the spatial spread of attention? Indeed, when we looked at our stimuli, we found that control objects tended to be further away from the main agent object at the moment when the simulation probe occurred (Figure S3A). Additionally, in the debrief survey, several participants mentioned focusing on the moving object as a strategy. To address this potential confound, we ran an analysis on a subset of trials where we matched the pixel-based Euclidean distance between the target object and the agent object (e.g., the red ball that will soon collide with the target object, but not the control object) and the control object and the agent object. Specifically, we only looked at trials where the distance between the center of the agent and the center of the target/control object was between 225 to 275 pixels, reducing the number of target and control object trials per participant from approximately 50 to between 10 and 20 trials. When matching for spatial distance in this way, we found a similar (in fact, numerically larger) effect as in our main result: participants’ detection accuracy for simulation probes that occurred on target objects (*M* = 55.11%) was significantly higher than that of control objects (*M* = 41.29%; *t*(27) = 3.26, *p* = .003, Cohen’s d = 0.63; Figure 5C). This effect was again evident in the majority of participants (17 out of 28; Figure 5D), and robust across the range of spatial distance values that were tested (Figure S3B).

We also investigated whether the particular letters we used as probes mattered for our effects. To our surprise, we observed that detection accuracy as well as the magnitude of the effect were impacted by the identity of the letter shown (see Figure S4A, B). Even though our main aggregate results comparing target and control objects are not impacted by this difference between the two letter probes, this result indicates that future studies should consider the detectability of letters when designing their experiments (Mueller & Weidemann, 2012).

### Confidence and response time also reflect simulated trajectory of objects

We also tested whether participants’ confidence and response times reflect any signature of simulated trajectories within an event. First, we investigated confidence in probe detection accuracy averaged across target and control probe trials within each participant and used a two-sided paired *t*-test to determine significance. As depicted in Figure S5A, participants’ confidence in their performance at detecting simulation probes appearing on target objects (*M* = 61.86) was higher than their confidence in detecting simulation probes appearing on control objects (*M* = 57.47; *t*(27) = 3.28, *p* = .003). Second, to analyze response time data, we considered participants who responded to at least one target and control trial, resulting in only 26 participants. Trending in the direction we expect, we found participants’ response times were numerically smaller on target objects (*M* = 1603.07 ms) relative to controls (*M* = 1635.37 ms; *t*(25) = -1.83, *p* = .078), but this relationship did not reach significance (Figure S5B). These results are consistent with our main detection accuracy findings, suggesting that objects along the simulated trajectory of the current event are prioritized.

### Connecting Experiments 1 and 2: Detection accuracy at event boundaries versus event middles

Thus far, the results from Experiment 2 have shown that physical events enable selective processing of objects that are on the future trajectory of the current physical event, relative to control objects. At a finer-level of granularity, the ability to detect a simulation probe may depend on when during the time course of an event it occurs, much like the time course of the detectability of temporal probes in Experiment 1. Because simulation should be tied to the current physical event, we may expect the detection accuracy of simulation probes to be higher during an event (event middle) compared to at an event boundary.

In Experiment 2, simulation probes were typically placed just before physical event boundaries, but sometimes, they appeared much earlier in time ***—*** i.e., during the middle of an event. This variability in the temporal placement of simulation probes gave us the opportunity to explore whether detection accuracy was modulated by temporal proximity to the event boundary. Experiment 1 demonstrated that temporal probe detection accuracy is at baseline levels around 600 ms before the event boundary, but substantially drops within 200 ms of the event boundary. Therefore, we labeled probes that occurred 600 ms or more before the event boundary ‘event middle simulation probes’ and probes that occurred within 200 ms ‘event boundary simulation probes.’ We first examined trials in which the simulation probe appeared on a target object (‘target trials’), averaging probe detection accuracy for event middle simulation probes vs. event boundary simulation probes within each participant and using a two-sided paired *t*-test to determine significance. Participants’ detection accuracy for target simulation probes that occurred at event boundaries (M = 47.17%) was significantly lower than for probes that occurred at event middles (M = 54.79%; t(27) = -2.70, p = .012, Cohen’s d = -0.52; Figure 6A). These results show that detection accuracy decreases for target simulation probes at event boundaries, relative to event middles, and are broadly consistent with the event cognition literature (Zacks et al., 2007). We emphasize that our findings are based on measuring spontaneously unfolding processes in visual perception, and underscore the importance of the temporal structure imposed by physical events on perception.

**Figure 6.**
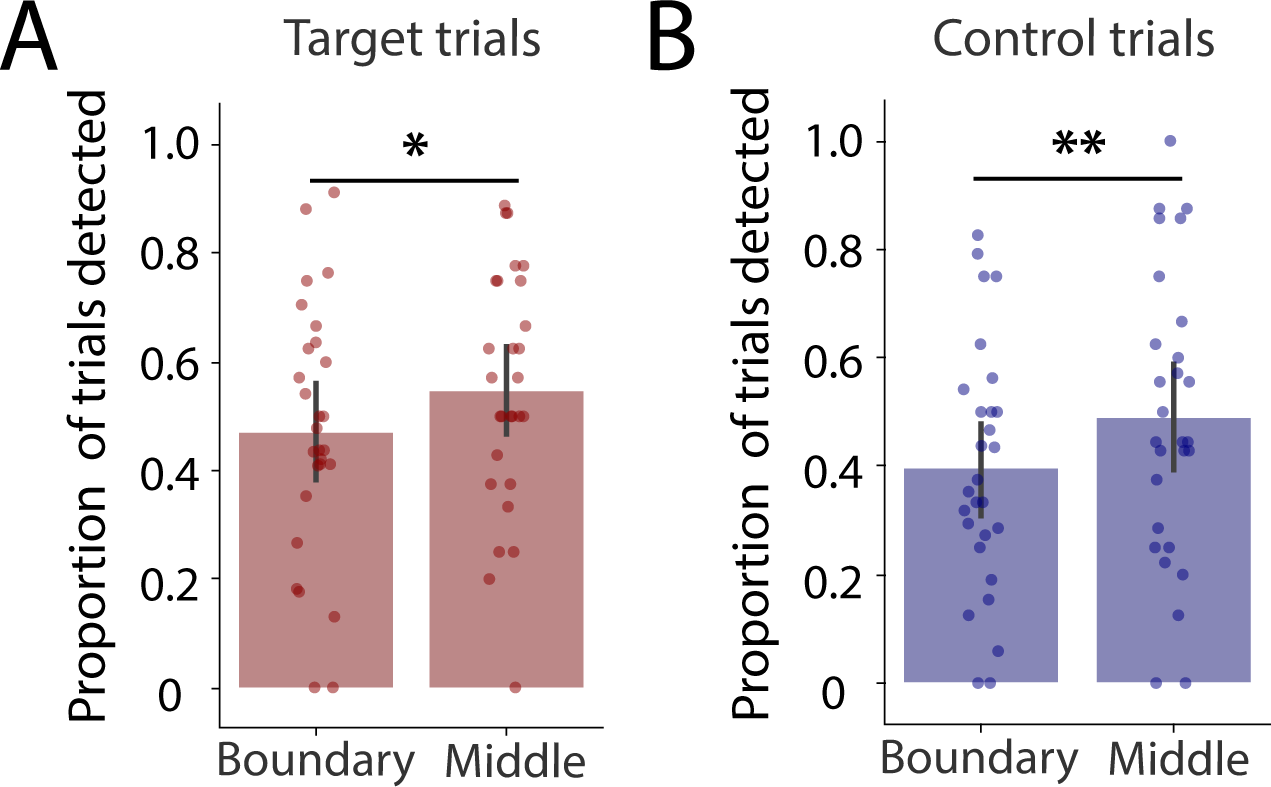
Simulation probe detection accuracy in Experiment 2 based on when during the physical event the probe appeared. (A) Target simulation probes were significantly more likely to be detected when they appeared at event middles (more than 600ms from an event boundary) compared to event boundaries (within 200 ms of an event boundary). Dots represent individual participant averages, and error bars reflect the 95% confidence interval derived from bootstrap resampling of the mean. (B) Control probes were also significantly more likely to be detected when they appeared at event middles compared to event boundaries. ** *p* < .01, * *p* < .05.

However, if the temporal-dependence of the ability to detect simulation probes is exclusively due to event-driven prediction (e.g., as assumed in Eisenberg et al., 2018), we may expect to not see a temporal-dependence for objects that are irrelevant to the ongoing event. To test this, we examined participants’ ability to detect probes that occurred on control objects (‘control trials’). We found that participants’ detection accuracy for control probes that occurred at event boundaries (M = 39.39%) was also significantly lower than for probes that occurred at event middles (M = 48.63%; t(27) = -2.98, p = .006, Cohen’s d = -0.57; Figure 6B). Thus, probe detection accuracy is enhanced at event middles not just for those objects that are relevant to the current physical event, but also for control objects that are in the vicinity of the target object. These results suggest that the decreased ability to detect simulation probes at event boundaries may reflect a more global effect of attention differences, instead of, or in addition to, event-driven prediction.

## General Discussion

In this paper, we conducted two performance-based tasks to gain insight into the spontaneous formation and contents of physical events in visual perception. Our first experiment confirmed that physical events are spontaneously segmented in vision, and allowed us to reveal the temporal profile of how such segmentation occurs during ongoing perception, at sub-second resolution and at the level of individual event-types. Our second experiment queried the contents of physical events based on the principle of prediction. We found that objects along the simulated trajectory of the current physical event have an attentional advantage over similar objects outside of such predicted trajectory. Together, these results provide an empirical basis for understanding physical events as units of visual perception during ongoing experience, starting to bridge the knowledge gap relative to our understanding of visual object representations and highlighting new avenues for exploration in intuitive physics.

We note that our study is not unique in the study of physical events in intuitive physics; several studies have used physical events, but typically in the form of pre-segmented singleton events (e.g., animations including a single event or interaction) which act as stimuli to study some other cognitive or perceptual phenomena (Gerstenberg et al., 2017, 2021; Schwettmann et al., 2019). As far as we can tell, physical events are not the ‘object’ of study in this work, i.e., they do not address the structure of physical events, including how they are segmented to begin with and their contents. Thus, our work makes an important advance in understanding how physical events impose temporal structure and influence the contents of intuitive physics, providing the scaffolding for these effects to be observed.

The task in Experiment 1 was inspired by work in audition and music cognition (Repp, 1992, 1998), which found that participants are less likely to detect musical phrase lengthenings when they are placed at boundaries. Transferring this method to the domain of vision allowed us to determine when event boundaries are perceived during a series of physical interactions, without informing participants to attend to physical event structure. Our results mirror that of Ji and Papafragou (2022), who found that participants are less likely and slower to detect visual disruptions at event boundaries compared to event middles (particularly for the so-called ‘bounded’ events, relative to ‘unbounded’ events, an important distinction made in the linguistic theories of events). In the current study, we tested not just event boundaries and middles, but also other timepoints in between, allowing us to quantify event segmentation in a temporally-precise manner. This approach provides a data-driven ‘behavioral signature’ of event boundary status that should be broadly useful across psychological and cognitive neuroscience studies of event perception. A key question left unanswered in the current study is *why* participants are worse at detecting temporal probes at physical event boundaries. One possibility is that these event boundaries simply correspond to moments of perceptual change (Newtson, Engquist, & Bois, 1977; Hard, Recchia, & Tversky, 2011). However, our analysis controlling for changes in pixels indicates that a purely low-level explanation is unlikely to account for this result. Rather, the lower temporal probe detection accuracy may reflect instantiation of a new event model, which likely consumes additional cognitive resources and might therefore impact task accuracy. Indeed, this fits with our results showing that the impact of an event boundaries on probe detection accuracy is not just at the moment of the boundary, but also during the lead up and follow up of the events. A related possibility is that event transitions ‘feel’ subjectively longer (therefore making the temporal probes not feel out of place), as suggested in time perception research: Time is judged to have lasted longer when multiple events occur (Faber & Gennari, 2015), and rhythms are reproduced as longer at event boundaries (Ongchoco, Yates, & Scholl, in press).

Infant cognition research has utilized physical event types to study how physical reasoning develops early in life. One surprising and consistent finding is that infants treat containment and occlusion differently (Baillargeon, 2004; Baillargeon et al., 2011; Hespos & Baillargeon, 2016; Lin et al., 2021). Paralleling this result, we found that detection accuracy showed temporally symmetrical decay and growth relative to containment events, but this pattern was temporally asymmetric for occlusion events. Similar to recent work exploring intuitions about magic tricks (Lewry et al., 2021), future research should systematically study parallels between intuitive physics within the adult visual system and its developmental trajectory.

Experiment 2 adapted and tested a key idea from the event cognition literature (Zacks et al., 2011) in visual perception of physical events: that events are defined by their ability to predict what is going to happen next. For physical events, prediction entails simulating the trajectory of a moving object and anticipating which new objects will interact with the moving object (a form of coarse simulations, or ‘looking ahead’). Participants were better at detecting probes that occurred on objects along the simulated trajectory of the current event. Physical events and simulation within them may therefore be a general principle that determines the contents of physical events, including the relevant features of objects (Strickland & Scholl, 2015), which objects themselves are relevant, and perhaps even inducing invisible ones into perception (Little & Firestone, 2021).

Although we describe this effect as the result of ‘look ahead’ simulations, we do not have eye-tracking data to say whether or not participants made predictive fixations towards to the target object during the task. Nonetheless, because target and control objects were probed equally often in the experiment, any predictive looking due to task or goals should have been equally likely for either object (Hayhoe & Ballard, 2005). An explanation that our results are due to distance to the moving object does not hold because our effects were still present (and numerically larger) when restricting our analyses to target/control objects that were at a similar distance to the moving object. Thus, we believe that these results reflect event-driven effects on object-based attention (Chen, 2012). Interestingly, while we replicated the finding that probes placed at event boundaries are more difficult to detect than probes placed at event middles (Huff et al., 2012), this effect was not specific to the target objects. Thus, the decreased ability to detect probes at event boundaries appears to be a more global impact of attention, rather than something that is specific to the ongoing event. These results underscore the importance of having fine-grained controls not only in terms of temporal aspects of a scene (e.g., as in Ji & Papafragou, 2022; Eisenberg et al., 2018), but also the object-level contents of what might be influenced by events.

There are several limitations to note in this study. First, there was a difference in the number of boundary versus non-boundary trials that were tested within each participant in Experiment 1. This was part of our initial design decision, given that we wanted to sample as many different timepoints in each video as possible, but likely impacted power (e.g., the analysis in Fig. 3D focusing on boundary trials). Second, despite interesting event-specific effects in Experiment 1, we did not have the power to break Experiment 2 data down into individual event types because of the smaller participant sample size and number of videos tested. Third, in Experiment 2, we only investigated attention allocation at time points near event boundaries or about one second away with a single control object. Further mapping out how attention is directed, and when it is heightened, during physical events will help uncover more of the internal structure of physical events.

Our work underscores the existence and structure of physical events in visual perception, which should inform computational accounts of intuitive physics (Bear et al., 2021; Piloto et al., 2022). An influential framework suggests that the mind builds runnable mental models via an ‘intuitive physics engine’ similar to probabilistic or approximate forms of physics engines from computer graphics (Battaglia et al., 2013). Recent work observed the implausibility of simulating the trajectories of all objects in a scene all the way to the end of their motions (Ludwin-Peery et al., 2021, Bass et al., 2021), but their solutions either considered a more heuristic-driven (and less simulation-like) basis for intuitive physics or suggested ad-hoc remedies. Our findings motivate an alternative for visual perception: minimally-expressed, selective simulations that are sufficient to explain and predict the trajectories of objects within their local causal contexts. Future computational work should implement such ‘event-driven intuitive physics engines’ to explore the mechanistic basis of physical events in perception.

## Acknowledgements

T.S.Y. was supported by National Science Foundation Graduate Research Fellowship, Grant/Award Number: DGE 1752134. I.Y. was supported by AFOSR grant # FA9550-22-1-0041. We are thankful to the participants and the CNCL lab for their helpful feedback on earlier iterations of this project. Thank you also to Winnie Chen for help in the creation of stimuli for Experiment 2.

## Author Contributions

**T.S.Y.**: Conceptualization, Formal analysis, Software, Writing – original draft, Writing – review and editing; **S.Y.**: Conceptualization, Software, Writing – review and editing; **I.Y.**: Conceptualization, Supervision, Writing – original draft, Writing – review and editing

## Data and code availability

De-identified data for Experiments 1 and 2, as well as the videos that were shown, are publicly accessible at https://osf.io/ma32p/. The code used to run the tasks is publicly accessible on Github (Experiment 1 temporal probe task: https://github.com/CNCLgithub/phys-env-psiturk/tree/temporal_probe; Experiment 1 event segmentation task: https://github.com/CNCLgithub/phys-env-psiturk/tree/event_seg; Experiment 2 spatial probe task: https://github.com/CNCLgithub/phys-env-psiturk/tree/spatial_probe). The code used to create the videos for both experiments is also publicly accessible on Github (https://github.com/CNCLgithub/physical_event_primitives) as is the code for the analyses (https://github.com/CNCLgithub/physical_events_analysis). There is a preregistration for Experiment 1 here https://aspredicted.org/blind.php?x=5ZF_D5T; Experiment 2 was not preregistered.

## Author Note

De-identified data for Experiments 1 and 2, as well as the videos that were shown, are publicly accessible at https://osf.io/ma32p. The code used to run the tasks is publicly accessible on Github (Experiment 1 temporal probe task: https://github.com/CNCLgithub/phys-env-psiturk/tree/temporal_probe; Experiment 1 event segmentation task: https://github.com/CNCLgithub/phys-env-psiturk/tree/event_seg; Experiment 2 spatial probe task: https://github.com/CNCLgithub/phys-env-psiturk/tree/spatial_probe). The code used to create the videos for both experiments is also publicly accessible on Github (https://github.com/CNCLgithub/physical_event_primitives) as is the code for the analyses (https://github.com/CNCLgithub/physical_events_analysis). There is a preregistration for Experiment 1 here https://aspredicted.org/blind.php?x=5ZF_D5T; Experiment 2 was not preregistered.

