## Supplemental Information for "Temporal segmentation and ‘look ahead’ simulation: Physical events structure visual perception of intuitive physics"

### Supplemental Material

#### Supplemental Tables

Temporal probe detection confidence by pixel change + boundary condition

| | Beta estimate | Std. error | *t*-statistic | *p*-value |
|---|---|---|---|---|
| Intercept | 73.952 | 1.248 | 59.246 | < .001 |
| Condition (boundary) | -6.974 | 0.933 | -7.472 | < .001 |
| Pixel change | 31.497 | 5.231 | 6.021 | < .001 |
| Participant random effect | 155.857 | 1.151 | - | - |

**Table S1.** Results from a general linear model predicting confidence from the amount of pixel change at the probe and the boundary condition (1 for boundary, 0 for non-boundary) across all participant trials (with participant as a random effect). The equation used in the model is shown as the table heading. The beta parameter, standard error, *t*-statistic, and *p*-values are given for each of the predictors.

Temporal probe detection response time by pixel change + boundary condition

| | Beta estimate | Std. error | *t*-statistic | *p*-value |
|---|---|---|---|---|
| Intercept | 1626.472 | 29.315 | 55.483 | < .001 |
| Condition (boundary) | 69.626 | 20.722 | 3.360 | .001 |
| Pixel change | -999.436 | 174.892 | -5.715 | < .001 |
| Participant random effect | 81280.833 | 44.601 | - | - |

**Table S2.** Results from a general linear model predicting response time (ms) from the amount of pixel change at the probe and the boundary condition (1 for boundary, 0 for non-boundary) across all participant trials (with participant as a random effect). Note that trials in which the participant did not respond with a key press are dropped from the analysis. The equation used in the model is shown as the table heading. The beta parameter, standard error, *t*-statistic, and *p*-values are given for each of the predictors.

#### Supplemental Figures

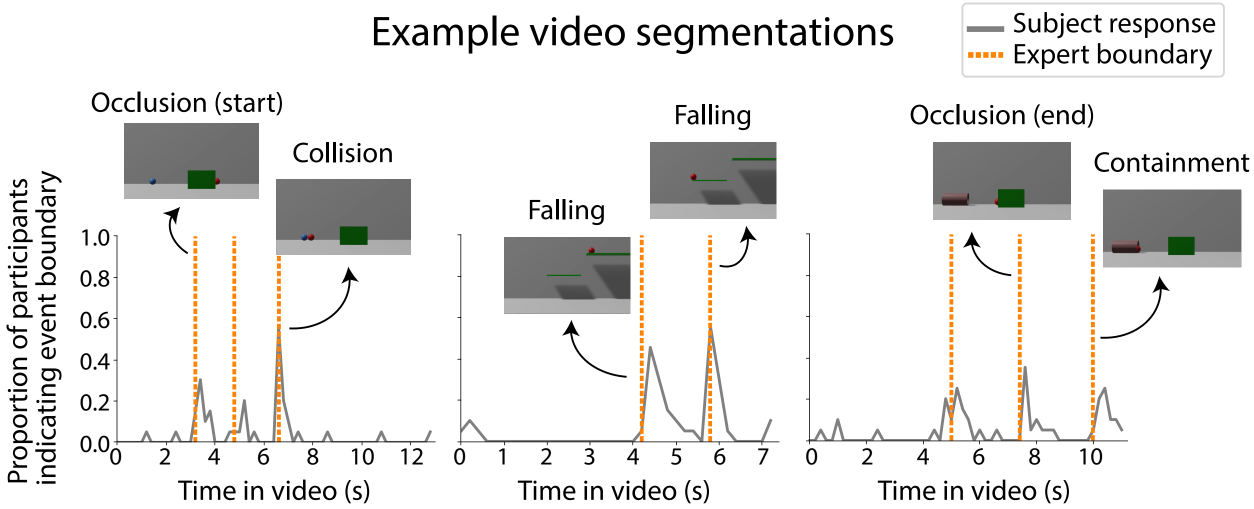


**Figure S1.** Example time courses of event segmentation by naive participants tasked with explicitly demarcating 'events' during videos containing different physical event types. Participants were allowed to indicate as many event boundaries as they saw fit, and could move forward and backward through the videos to precisely indicate their event boundary judgment. Gray lines indicate the proportion of event segmentation participants (out of 20) who indicated that there was an event boundary at that time point. Dashed orange lines indicate the expert-determined event boundaries used in our main analyses. Screenshots of expert boundaries are inset.

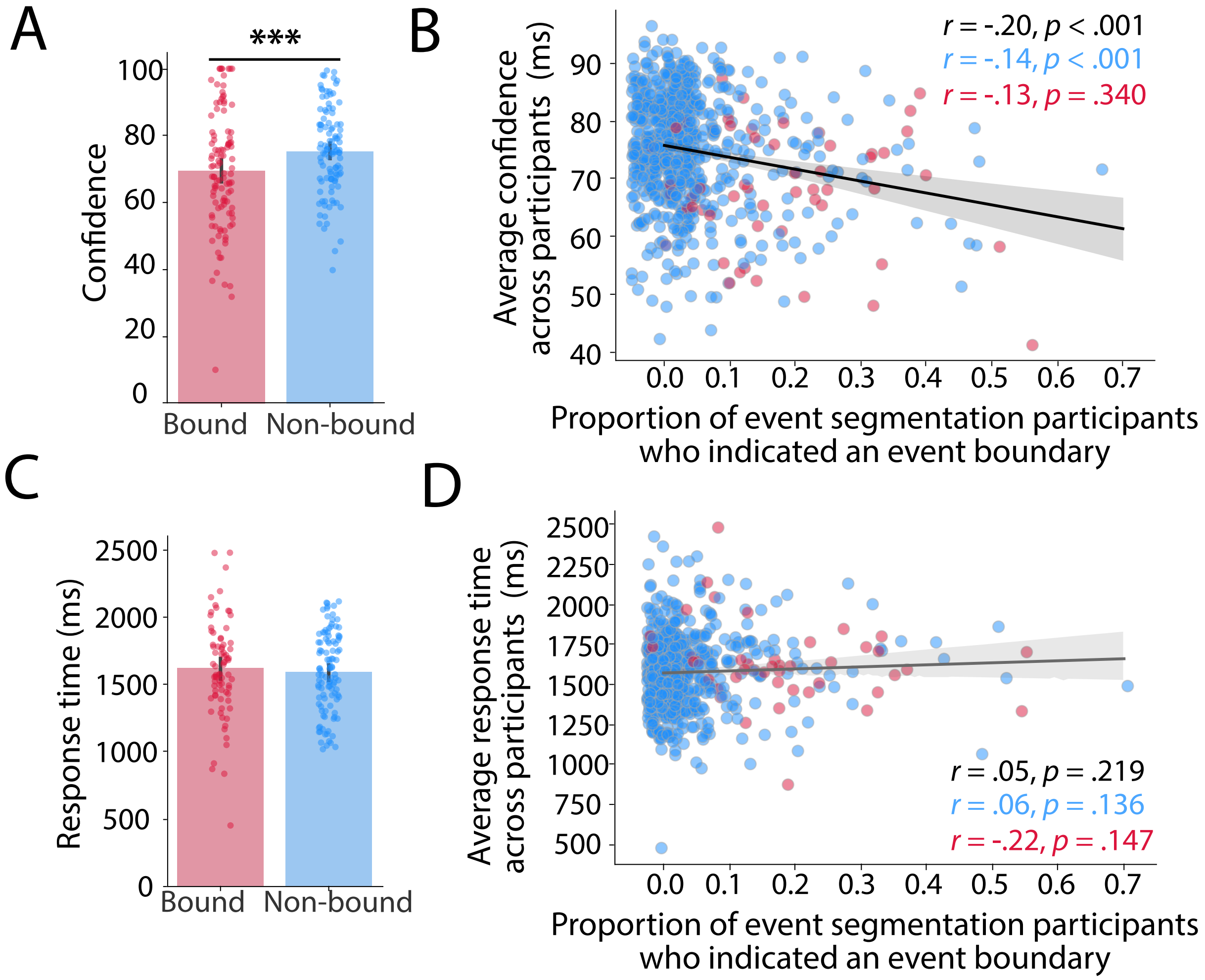


**Figure S2.** Temporal probe detection confidence, and response times from Experiment 1, broken down by participant. (A) Confidence in participants' probe detection accuracy was higher on non-boundary trials compared to boundary trials. Dots represent individual participant averages, and error bars reflect the 95% confidence interval derived from bootstrap resampling of the mean. (B) However, confidence was not significantly related to the proportion of event segmentation participants who indicated an event boundary around that time point. (E) For trials in which a probe was detected, response times for boundary probes ($M$ = 1616.84 ms) were numerically larger than response times for non-boundary probes ($M$ = 1586.36 ms; $t(77) = 1.92$, $p = .059$). (F) Response time was not associated with the proportion of event segmentation participants who indicated an event boundary around that time point when considering all probes ($r = .05$, $p = .219$), only non-boundary probes (blue dots; $r = .06$, $p = .136$), or only boundary probes (red dots; $r = -.22$, $p = .147$). Dots represent individual participant averages, and error bars reflect the 95% confidence interval derived from bootstrap resampling of the mean. *** $p < .001$.

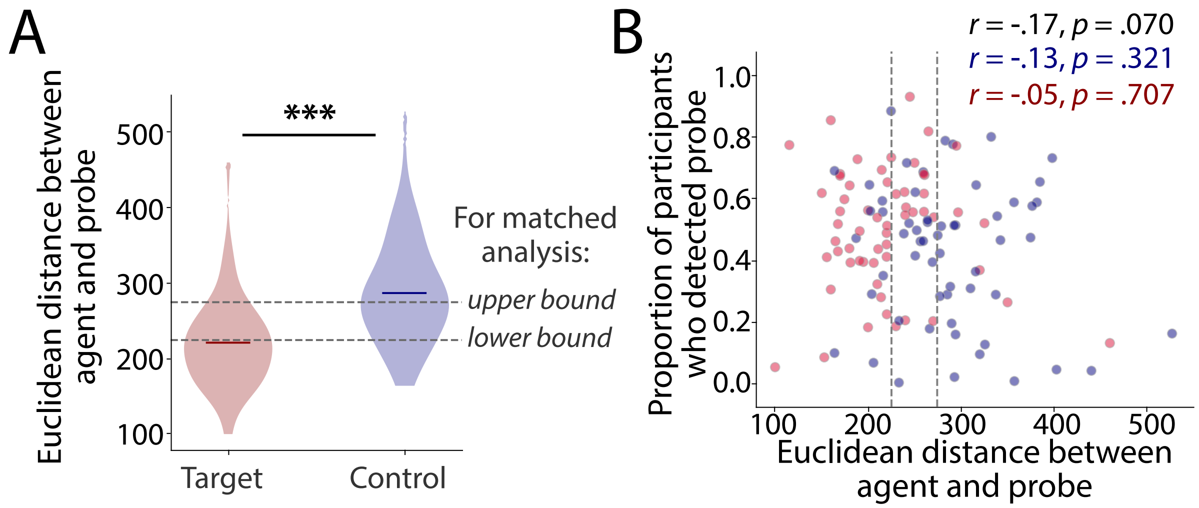


**Figure S3.** (A) Euclidean distance in pixels between the center of the agent object (e.g., the red ball that is about to roll behind an occluder) and the center of the flashing simulation probe across all possible trials (58 each). Target objects tended to be closer to the agent object than to control objects. (B) Detection accuracy did not significantly vary by the euclidean distance between the agent and the probe. Dashed gray lines indicate the distances used for the matched-distance analysis. *** $p < .001$

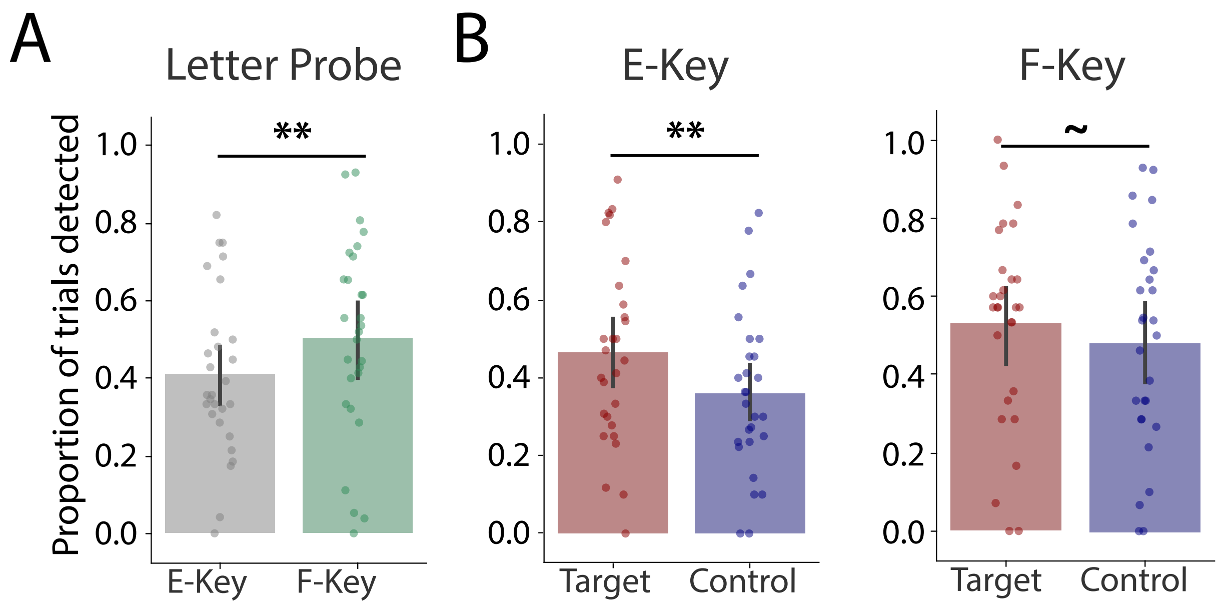


**Figure S4.** Breakdown of simulation probe detection accuracy by letter type (‘E’ or ‘F’). (A) Probes that flashed the letter ‘F’ were more easily detected than probes that flashed the letter ‘E’. (B) For ‘E’ trials, participants were more likely to detect probes on target objects compared to control objects. For ‘F’ trials, this difference was not significant, although there was still a trend towards better detection of target objects. ** $p < .01$, ~ $p < .10$

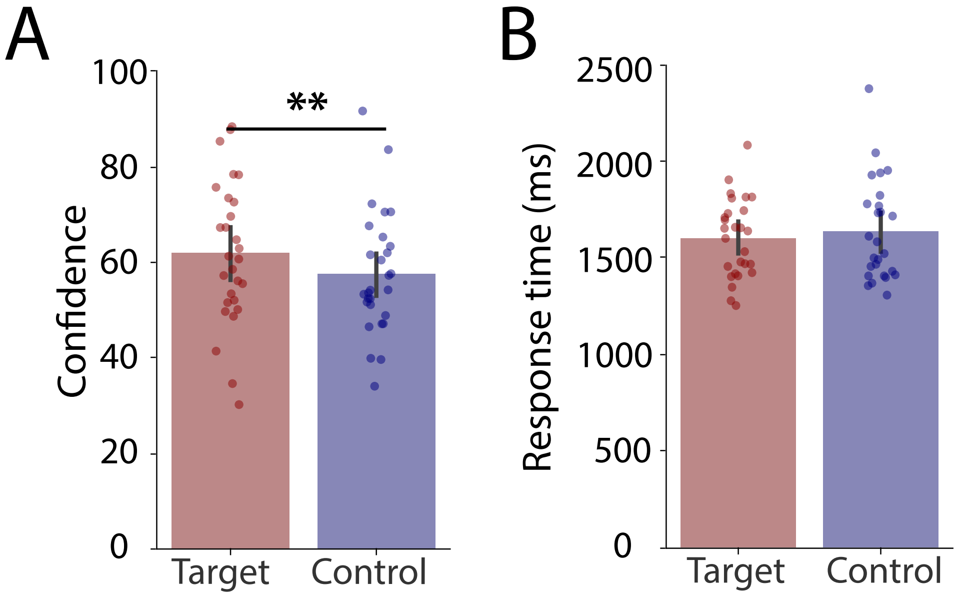


**Figure S5.** Simulation probe detection confidence, and response times from Experiment 2, broken down by participant. (A) Confidence in participants' probe detection accuracy was higher on target trials compared to control trials. Dots represent individual participant averages, and error bars reflect the 95% confidence interval derived from bootstrap resampling of the mean. (B) For trials in which a probe was detected, response times for target objects ($M$ = 1603.07 ms) were numerically smaller than response times for probes on control objects ($M$ = 1635.37 ms; $t(25) = -1.83$, $p = .078$). ** $p < .01$.
